# From root to tips: sporulation evolution and specialization in *Bacillus subtilis* and the intestinal pathogen *Clostridioides difficile*

**DOI:** 10.1101/473793

**Authors:** Paula Ramos-Silva, Mónica Serrano, Adriano O. Henriques

## Abstract

Bacteria of the Firmicutes phylum are able to enter a developmental pathway that culminates with the formation of a highly resistant, dormant spore. Spores allow environmental persistence, dissemination and for pathogens, are infection vehicles. In both the model *Bacillus subtilis*, an aerobic species, and in the intestinal pathogen *Clostridioides difficile*, an obligate anaerobe, sporulation mobilizes hundreds of genes. Their expression is coordinated between the forespore and the mother cell, the two cells that participate in the process, and is kept in close register with the course of morphogenesis. The evolutionary mechanisms by which sporulation emerged and evolved in these two species, and more broadly across Firmicutes, remain largely unknown. Here, we trace the origin and evolution of sporulation. Using the genes involved in the process in *B. subtilis* and *C. difficile*, and estimating their gain-loss dynamics in a comprehensive bacterial macro-evolutionary framework we show that sporulation evolution was driven by two major gene gain events, the first at the base of the Firmicutes and the second at the base of the *B. subtilis* group and within the Peptostreptococcaceae family, which includes *C. difficile*. We also show that early and late sporulation regulons have been co-evolving and that sporulation genes entail greater innovation in *B. subtilis* with many Bacilli-lineage restricted genes. In contrast, *C. difficile* more often recruits new sporulation genes by horizontal gene transfer, which reflects both its highly mobile genome, the complexity of the gut microbiota and an adjustment of sporulation to this particular ecosystem.

## Introduction

Bacterial endospores (spores for simplicity) are among the most resilient cells known to us. Because of their resilience to extremes of several physical and chemical conditions (Setlow 2014), spores can spread and persist in the environment for long periods of time and in a variety of ecosystems (Nicholson et al. 2000), from the human gut (Hong et al. 2009; Browne et al. 2016) to the deep sea (Urios 2004; Dick et al. 2006; Fang et al. 2017). Spores can also have a profound impact on human health (Yano et al. 2015; Browne et al. 2016). Spore-formers can either be symbiotic (Angert and Losick 1998; Flint et al. 2005), pathogenic (Abt et al. 2016) or free-living (Ramos-Silva et al. 2015; Fang et al. 2017). Spores have a core compartment that encloses the DNA delimited by a membrane and surrounded by a thin layer of peptidoglycan that will serve as the cell wall of the new cell that emerges from spore germination. A second layer of modified peptidoglycan essential for dormancy, the cortex, is surrounded by several proteinaceous layers (Henriques and Moran 2007). In *Bacillus subtilis* a multilayered protein coat forms the spore surface (Lai et al. 2003; Kim et al. 2006); in other species, such as *Bacillus anthracis* and *Clostridioides difficile* the spore is further enclosed in a glycosylated exosporium (Redmond 2004; Permpoonpattana et al. 2011; Thompson et al. 2012).

Sporulation usually begins with the formation of a polar septum that leads to an asymmetric cell division resulting in two cells, the forespore (or future spore) and the mother cell, each with an identical copy of the chromosome but expressing different genetic programs. At an intermediate stage, the forespore is engulfed by the mother cell, and becomes isolated from the external medium. At later stages, while gene expression in the forespore prepares this cell for dormancy, the mother cell drives the assembly of the cortex, coat and exosporium, and eventually triggers its own lysis to release the mature spore (Higgins and Dworkin 2012). Entry in sporulation is triggered by the activation of the transcriptional regulator Spo0A (Fujita and Losick 2005). Spo0A is conserved among spore-formers and its presence is often used as an indicator of sporulation ability (Galperin 2013; Filippidou et al. 2016). The activity of Spo0A is required for the division that takes place at the onset of sporulation. Following division, gene expression in the forespore and the terminal mother cell is sequentially activated by cell type-specific RNA polymerase sigma factors (Losick and Stragier 1992; Stragier and Losick 1996; Kroos and Yu 2000; Rudner and Losick 2001): prior to engulfment completion, gene expression is governed by σ^F^ in the forespore and by σ^E^ in the mother cell; following engulfment completion, σ^F^ is replaced by σ^G^ and σ^E^ is replaced by σ^K^. Like Spo0A, sporulation sigma factors are also conserved among spore-formers (de Hoon et al. 2010; Galperin et al. 2012; Abecasis et al. 2013). The identification of the genes whose transcription is directly controlled by σ^F^, σ^E^, σ^G^ and σ^K^ was initially made in *Bacillus subtilis*, an aerobic organism, representative of the class Bacilli, which is still the best characterized spore-former (Fawcett et al. 2000; Eichenberger et al. 2003; Wang et al. 2006; Meisner et al. 2008; Mäder et al. 2012). *Clostridium difficile*, later re-classified as *Clostridioides difficile*, (Lawson et al. 2016), is a nosocomial pathogen and the main cause of intestinal diseases linked to antibiotic therapy worldwide (Bauer et al. 2011; Lessa et al. 2015; Abt et al. 2016). As an obligate anaerobe, *C. difficile* relies on its sporulation ability to spread horizontally between hosts via fecal and oral transmission (Lawley et al. 2009; Janoir et al. 2013). This has motivated a number of studies on the control of sporulation and *C. difficile* has emerged in recent years as the anaerobic model for the process, being currently the best characterized species within the Clostridia (Paredes-Sabja et al. 2014). Importantly, several studies have been directed at the identification of the genes under the control of the cell type-specific sigma factors (Fimlaid et al. 2013; Saujet et al. 2013), and several genes have been characterized, highlighting conservancy of function of the main regulatory proteins, but also major differences in the control of sporulation relative to the model *B. subtilis* (Fimlaid et al. 2015; Fimlaid and Shen 2015; Pishdadian et al. 2015; Serrano et al. 2016).

Based on the extensive knowledge of sporulation in *B. subtilis*, previous comparative genomics studies have led to the identification of a core (*i.e.* genes conserved in all spore-formers), a signature (present in all spore-formers but absent in non-spore-forming lineages) and Bacilli-specific sporulation genes (Onyenwoke et al. 2004; Galperin et al. 2012; Abecasis et al. 2013) as well as to elucidate how conservation patterns are related to gene functional roles (de Hoon et al. 2010).

Although restricted to Firmicutes, sporulation is a highly diversified process within the phylum (Angert and Losick 1998; Flint et al. 2005; Galperin 2013; Yutin and Galperin 2013; Hutchison et al. 2014). Still, the evolutionary mechanisms driving this diversity remain largely unknown. Here we address this question by comparative genomics of the *B. subtilis* sporulation genes together with the most recent set of sporulation genes from *C. difficile.* Special focus is given to genes whose expression is dependent on the cell type-specific sigma factors σ^F^, σ^E^, σ^G^ and σ^K^. We show that the σ regulons are highly divergent between the two species, which share only a small fraction of the genes. We then map the presence/absence of both sets of sporulation genes in a solid and comprehensive bacterial macro-evolutionary framework in order to estimate the origin and evolution of the sporulation gene families, from an ancient bacterial ancestor to the contemporary *B. subtilis* and *C. difficile.* Our analyses show common evolutionary patterns in the lineages of the two species, with two major gene gain events having expanded the cell type-specific regulons. Moreover, per each gain event, the cell type-specific regulons appear to grow in the same size-proportions, suggesting that sporulation stages have been co-evolving along the bacterial phylogeny, whereas multiple independent gene losses emerge in some branches of the tree for both Bacilli and Clostridia. We also show that in the evolution of *B. subtilis*, gene gains more often involve the emergence of taxonomically restricted genes (TRGs) than in *C. difficile*, where more sporulation genes have orthologous in other bacteria and were likely acquired by horizontal gene transfers (HGTs). Still, the two mechanisms are expected to occur to a greater or lesser extent in both lineages and are fully demonstrated here for the genes coding the exosporium protein CdeA in *C. difficile* (Díaz-González et al. 2015) and SpoIIIAF, which is a component of the SpoIIIA-SpoIIQ sporulation-essential secretion system conserved across spore formers (Meisner et al. 2008; Fimlaid et al. 2015; Morlot and Rodrigues 2018). Lastly, our analysis supports the hypothesis that sporulation emerged at the base of Firmicutes, 2.5 billion years ago, with a first major gene gain event in response to oxygen appearance. This major evolutionary step was followed by a series of intermediate gains that culminated with a second major gene gain event corresponding to the specialization of this developmental program and its adjustment to particular ecosystems.

## Results

### A small core and large diversity of sporulation genes between *Bacillus subtilis* and *Clostridioides difficile*

The species *Bacillus subtilis* is by far the best characterized spore-former in terms of sporulation genes and mechanisms (Fawcett et al. 2000; Eichenberger et al. 2003; Wang et al. 2006; Mäder et al. 2012). *Clostridioides difficile* is the second most-studied species and the best characterized within the class Clostridia (Fimlaid et al. 2013; Saujet et al. 2013; Paredes-Sabja et al. 2014; Pishdadian et al. 2015). For this study we have manually compiled 734 sporulation genes for *B. subtilis* strain 168 (Table S1, BSU) and 308 sporulation genes from the *C. difficile* strain 630 (Table S1, CD). Each gene is under the direct control of one or more sporulation-specific sigma factors and thus part of at least one sporulation regulon. The regulons have different sizes (Table 1): **σ**^E^ is the largest followed by **σ**^G^ and **σ**^K^ (of similar sizes) and **σ**^F^. Genes involved in sporulation and whose expression depends on Spo0A have only been thoroughly identified for *B. subtilis* (Fawcett et al. 2000).

**Table 1.** Number of sporulation genes collected in the literature per regulon and described in Table S1.

| **Sporulation stage** | **Expression** | **Regulon** | **# Genes**<br>***B. subtilis* 168** | **# Genes**<br>***C. difficile* 630** |
| --- | --- | --- | --- | --- |
| Initiation | Mother cell | Spo0A | 65 | - |
| Engulfment | Forespore | $\sigma^F$ | 81 | 39 |
| Engulfment | Mother cell | $\sigma^E$ | 335 | 204 |
| Maturation | Forespore | $\sigma^G$ | 156 | 60 |
| Maturation | Mother cell | $\sigma^K$ | 151 | 56 |

Homology mapping between *B. subtilis* and *C. difficile* genomes was carried out using a bi-directional BLASTP approach, followed by selection of sporulation gene clusters by regulon and presence/absence analysis of orthologous genes in both species. In Figure 1 we show that only a small fraction of sporulation genes have orthologous in the two species. Some of these genes are present in the same regulon (Figure 1, orange, Table S2): 2 genes in **σ**^F^, 29 in **σ**^E^, 11 in **σ**^G^, and 5 in **σ**^K^ while less than 1 % belongs to different regulons (yellow, Figure 1, Table S3). A larger proportion of genes corresponds to those present in both genomes but only reported to be involved in sporulation by one species (Table S4). These proportions vary depending on the regulon, in *B. subtilis* from 10 to 19% and in *C. difficile* from 12 to 35 %. Finally, the majority of sporulation genes have no candidate orthologous in the two species (Table S5) (> 70% for BSU, > 50% for CD), highlighting the great genetic diversity of the sporulation process. In order to elucidate the emergence of this diversity in a larger context of bacterial evolution we extended the analysis of presence/absence of sporulation genes to 258 bacterial genomes (Figure S1, Table S6).

**Fig 1.**
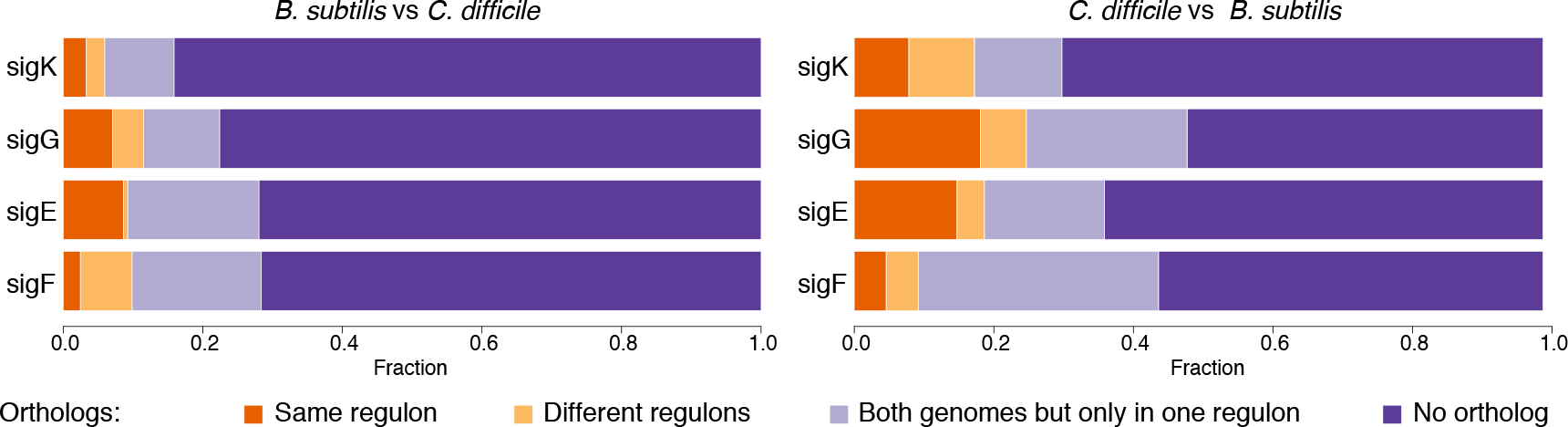
Orthology mapping between *B. subtilis* and *C. difficile* for genes under the control of sporulation sigma factors. The fraction of genes with orthologous in the same regulon is marked in orange. The fraction of genes with orthologous in different sporulation regulons is in yellow. The fraction of genes without orthologous in the regulons but with orthologous in the genome is in light purple. Fraction of genes without orthologous is in dark purple.

### Distribution of sporulation genes across bacteria reveals a small core of conserved genes and lineage-specificity

Presence/absence of sporulation genes was mapped across 258 bacterial genomes (Figure S1, Table S6) using an optimized orthology mapping approach (one *vs.* all and all *vs.* one) applied twice: first using *B. subtilis* 168 as reference, second using *C. difficile* 630 (Figure S2, Figure S3). The results were represented in super-matrices of presence (1 or more) and absence (0) of genes and used to calculate the distribution of sporulation genes across bacterial genomes (Figure 2). Both distributions show that only a small proportion of sporulation genes are conserved in the bacterial genomes (Figure 2A). A unimodal distribution is observed for sporulation genes from *B. subtilis* (D = 0.024757, p-value = 0.3179) with a peak around the 20%, whereas for *C. difficile* sporulation genes the distribution is bimodal (D = 0.05604, p-value = 7.262e-06) with high densities around 20% and a secondary high-density peak at 42% (Figure 2B). These results suggest that *B. subtilis* has more taxonomically restricted sporulation genes than *C. difficile*, whose genes appear more frequently in the bacterial genomic dataset. To better elucidate the causes underlying these differences we conducted an evolutionary analysis of sporulation gene families by posterior probabilities in a phylogenetic birth-death model.

**Fig 2.**
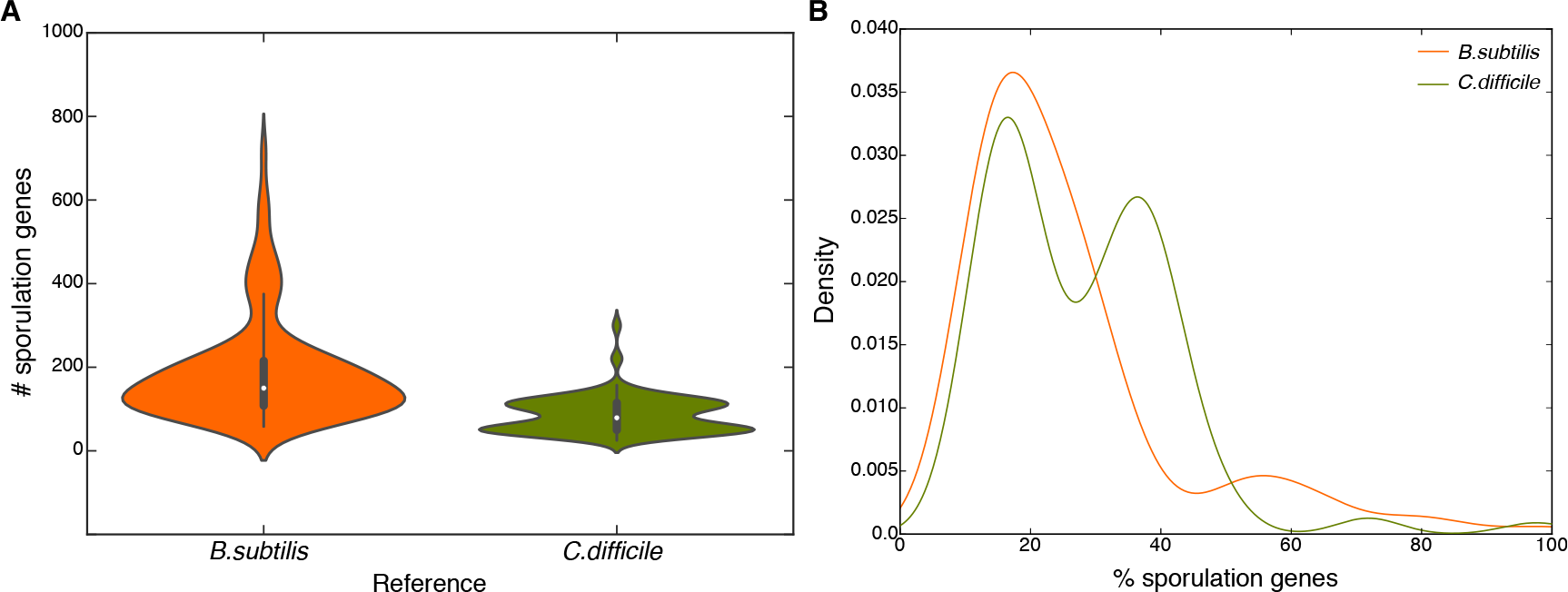
Distribution of sporulation genes from *B. subtilis* and *C. difficile* across the 258 bacterial genomes sampled from NCBI. Distributions are represented in (A) Absolute number of genes and (B) Relative percentages. Distributions were tested for unimodality using the Hartigan's Dip test in R (D = 0.024757, p-value = 0.3179 for *B. subtilis*; D = 0.05604, p-value = 7.262e-06 for *C. difficile*).

### A bacterial evolutionary backbone

Prior to the evolutionary analysis of sporulation gene families we started by building a solid and comprehensive bacterial phylogeny based on 70 marker genes conserved across the genomic dataset (Figure S1). Overall, the resulting tree topology (Figure 3, Figure S4) is similar to recent macroevolutionary bacterial trees (Hug et al. 2016; Marin et al. 2017). The phyla Dictyoglomi, Thermotogae, Aquificae, Synergistetes, and Deinococcus-Thermus form an early diverging clade. The second earliest split in the tree corresponds to the divergence of the taxon coined as hydrobacteria (which includes Proteobacteria, Bacteroidetes, Spirochaetes, Fusobacteria) and terrabacteria (which includes Actinobacteria, Cyanobacteria and the Firmicutes), estimated to have occurred between 2.83-3.54 billion years ago (Ga) (Battistuzzi et al. 2004; Battistuzzi and Hedges 2009). Within terrabacteria, Firmicutes (monoderms, low GC, mostly gram-positive) is an early diverging phylum currently divided into seven classes: Bacilli, Clostridia, Negativicutes, Erysipelotrichia, Limnochordia, Thermolitobacteria and Tissierellia (source: NCBI taxonomy). In our study we included genomes from four of these classes. The major split is between Bacilli (n=77) and Clostridia (n=95) but other groups elevated to the class level emerge within Clostridia, namely, the Negativicutes (n= 11) coding for diderm bacteria with a gram-negative-type cell envelope and Erysipelotrichia (n=1). Other two spore-formers (Figure S1), *Symbiobacterium thermophilum* and *Thermaerobacter marianensis* (gram-positive, GC > 60%) appear as a distinct lineage from the Firmicutes and have been suggested to be an intermediate phylum between the high-GC (Actinobacteria) and the low GC (Firmicutes) groups (Ueda et al. 2004; Han et al. 2010). Here, however, they remain as Firmicutes.

**Fig 3.**
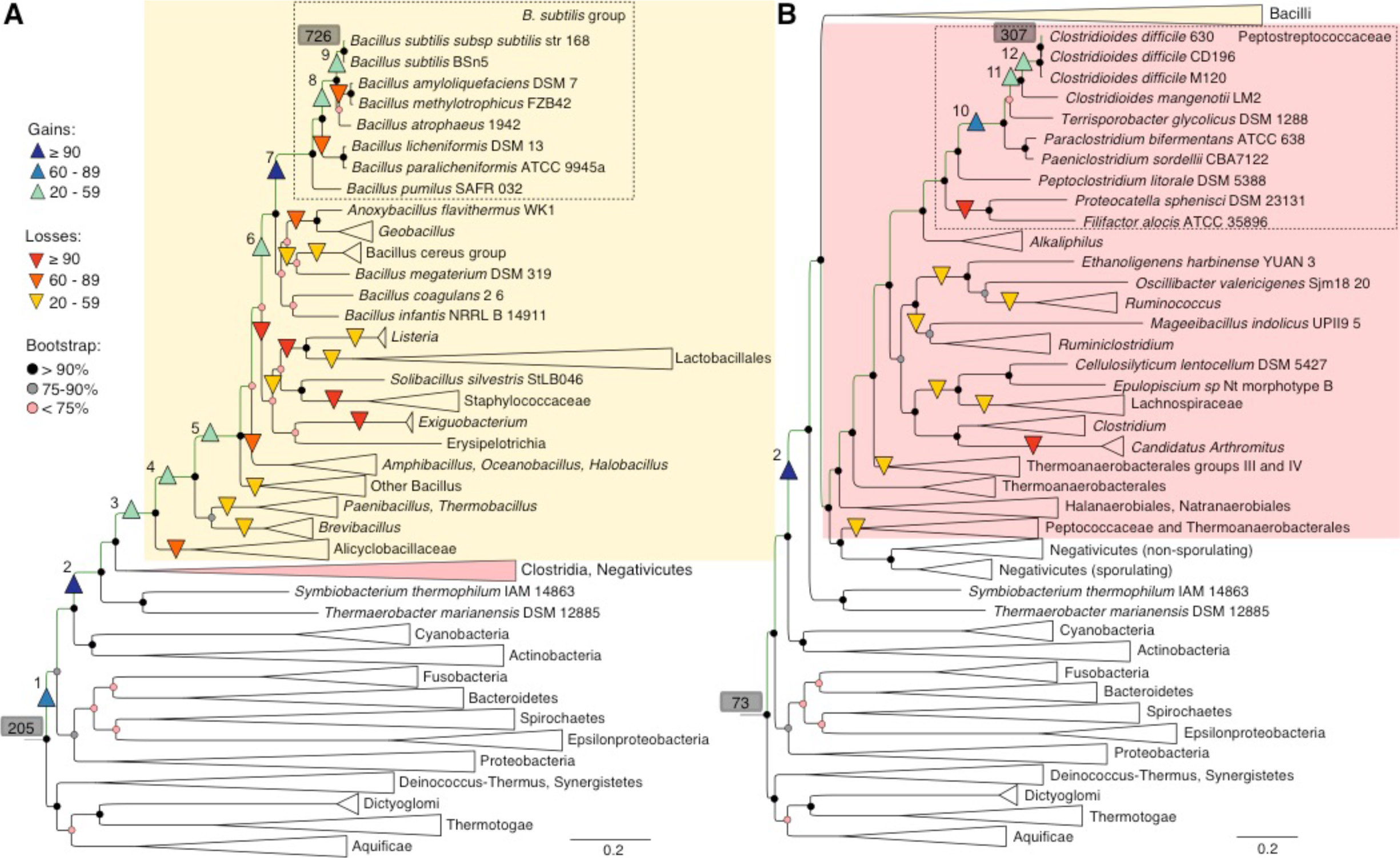
Sporulation gene gain and loss events across the bacterial phylogeny for (A) Bacilli and (B) Clostridia. The total number of sporulation genes present at the root and at the referential tips is highlighted in grey. Gain events are numbered from 1 to 12. Multigene maximum likelihood (RAxML) tree inferred from an alignment of 70 orthologs and corresponding bootstraps measures in %. The PROTGAMMALG evolutionary model was used to infer the tree with branch support estimated with 100 bootstrap replicates.

### Two major gene gain events drive sporulation evolution in *Bacillus subtilis* and *Clostridioides difficile*

Super matrices of presence/absence of sporulation genes (0,1, >1) sized 726×258 and 307×258 from references *B. subtilis* 168 and *C. difficile* 630, respectively, were used in an evolutionary analysis of sporulation genes across the bacterial phylogeny (Tables S7-S8, Figure 3). We first focused on the gene gain events, occurred from the root to the terminal branches of *B. subtilis* (Figure 3A) and *C. difficile* (Figure 3B). Our approach estimates that throughout evolution of both species, sporulation genes were acquired in two major evolutionary steps. The first major gene gain event occurred at the base of the Firmicutes, comprising at least 166 genes (branch 2, Figure 3, Table S9), including those coding for Spo0A and the four cell type-specific sigma factors, as well as other core sporulation genes, conserved among members of the phylum (Tables S9-S10). The second major gain event occurred at terminal branches, at the base of the *B. subtilis* group (Berkeley et al. 2002) (Figure 3A, +90 genes, branch 7, Figure 3A) as well as within members of the family Peptostreptococcaceae (Clostridium cluster XI) (Figure 3B, +73 genes, branch 10), at the branch that predates the split between the genera Clostridioides, Paeniclostridium, Paraclostridium and Terrisporobacter (Galperin et al. 2016). Intermediate gains also occurred in the evolutionary path of *B. subtilis* (branches 1, 3 to 6, 8 and 9) and *C. difficile*, in this case involving two additional gain events on branches 11 (+21 genes) and 12 (+39 genes) (Figure 3).

### Extensive gene loss in spore-forming and asporogenous lineages

Besides gene gains, also significant losses have occurred in certain Bacilli and Clostridia lineages. In Bacilli, extensive gene loss (Figure 3A, >=90, red triangles) was predicted for the branch ancestral to the asporogenous groups *Listeria* and Lactobacillales, *Solibacillus* and Staphylococcaceae, *Exiguobacterium* and Erysipelotrichia. In each of these groups, succeeding gene losses have occurred for Staphylococcaceae, *Exiguobaterium*, *Listeria* and Lactobacillales. In addition, multiple gene losses arose independently in other groups, including spore-formers, for *e.g.*, at the base of Alicyclobacillaceae, of the *Geobacillus* and *Anoxybacillus* genera and in some other *Bacillus* species (Figure 3A, orange triangles). In Clostridia, extensive gene losses have occurred independently for *Candidatus Arthromitus* and at the branch preceding the split of the genus *Filifactor* and *Proteocatella* (Figure 3B, red triangles). Other significant losses are predicted for the branches preceding the lineages Peptococcaceae and Thermoanaerobacterales groups III and IV, Thermoanaerobacterales, Lachnospiraceae, *Ruminiclostridium* and *Mageeibacillus*, and *Ruminococcus* (Figure 3B, yellow triangles).

Significant gains and losses were also predicted for some terminal branches of Bacilli and Clostridia species/strains (Table S7-S8). However, gain/loss expectations on terminal branches are highly prone to error because false positives/negatives, or lower quality genomes, have greater impact on the total estimates. Thus, only events occurring on internal branches were considered for discussion and presented in Figure 3.

### Co-evolution of early and late sporulation stages

Sporulation genes belong to at least one cell type-specific regulon and thus it is possible to explore the effect of gene gain events in the size of the regulons as a function of the evolutionary steps. Here we have focused on the gain events occurring throughout the evolutionary paths of *B. subtilis* and *C. difficile*, highlighted in green in Figure 3 (branches numbered from 1 to 12). The general pattern for *B. subtilis* (Figure 4A) and *C. difficile* (Figure 4B) is that regulons are growing in similar proportions. Larger increases in the size of the regulons corresponded to the major gene gain events highlighted in Figure 3: at the base of Firmicutes (branch 2), at the base of the *B. subtilis* group (branch 7) and within the family Peptostreptococcaceae (branch 10). In these branches we could also observe slightly larger increases for the regulons controlling late sporulation stages, **σ**^G^ (branch 2 and 10) and **σ**^K^ (branch 7) (Figure 4). Still these differences are small and do not affect the general pattern in which sporulation regulons seem to be co-evolving with gene gain dynamics until reaching their current size.

**Fig 4.**
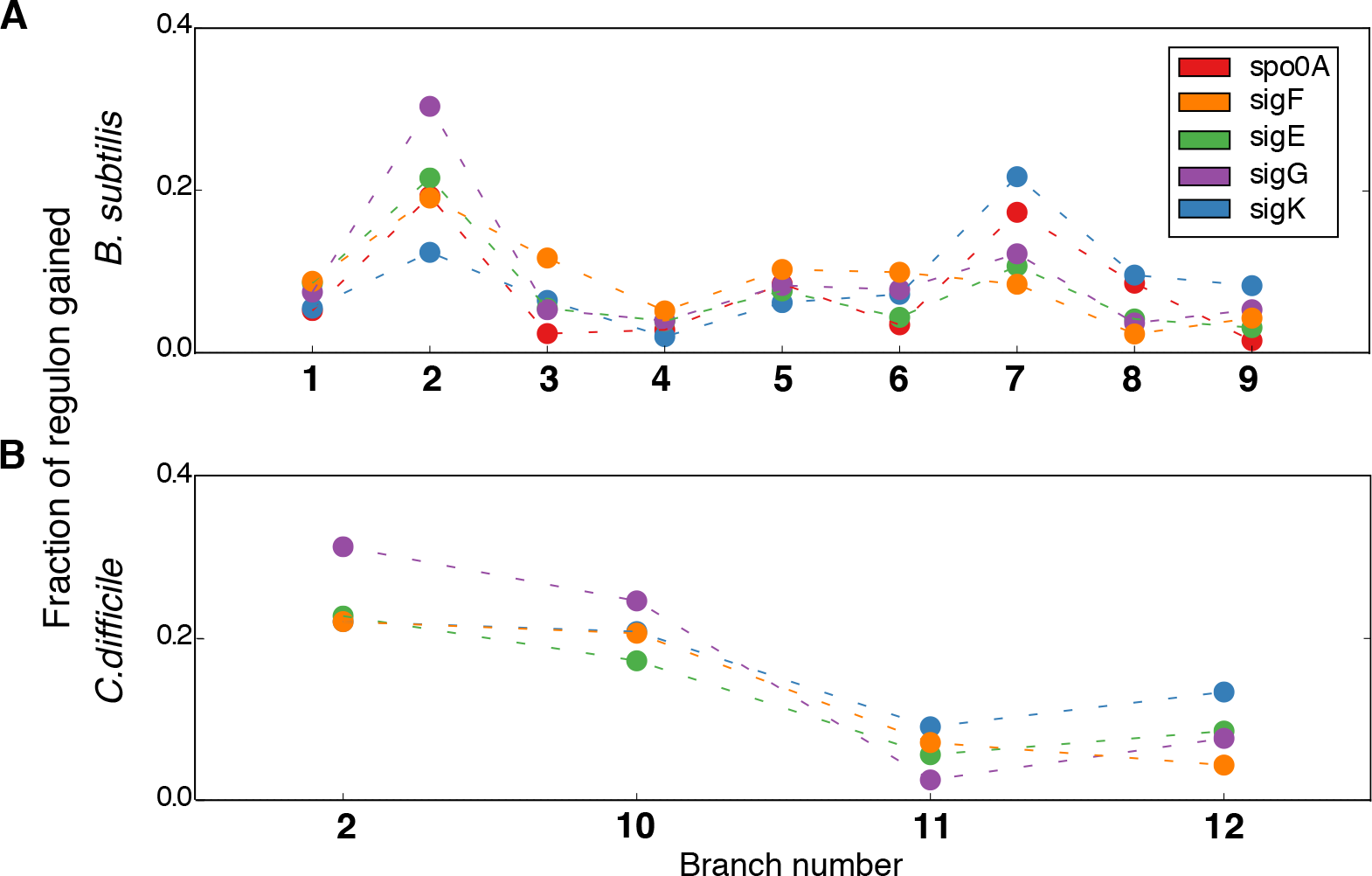
Increase in size of sporulation regulons represented from root to tips for (A) *B. subtilis* and (B) *C. difficile*. Sporulation regulons have diverse sizes being σ^**E**^the largest regulon and also the one acquiring the largest number of genes in every evolutionary step. To suppress this effect the gains per regulon were normalized by size.

### Sporulation genes: a landmark of Firmicutes

Evolutionary analyses of sporulation genes from *B. subtilis* and *C. difficile* estimate that 28,2% and 23,8% of their machinery was already present at the last bacterial ancestor (Figure 3, Tables S9-S10). These genes are conserved across early diverging bacterial clades and may have been co-opted for sporulation. Subsequent gene gains occurred later in the bacterial phylogeny, at different taxonomic levels and included TRGs, HGTs and duplications as illustrated in Figure 5. In particular, the first major gain event of sporulation genes (Figure 3, branch 2), which correlates with the emergence of the Firmicutes as a phylum, involved a gain of at least 166 genes, from which the vast majority, 80-90%, are TRGs, *i.e.* only found in Firmicutes members (Figure 5, Tables S9-S10). A much smaller amount of gene gains are estimated to be derived by HGT, 10,4-14,3% (Tables S9-S10), and having orthologous in other lineages outside the Firmicutes.

**Fig 5.**
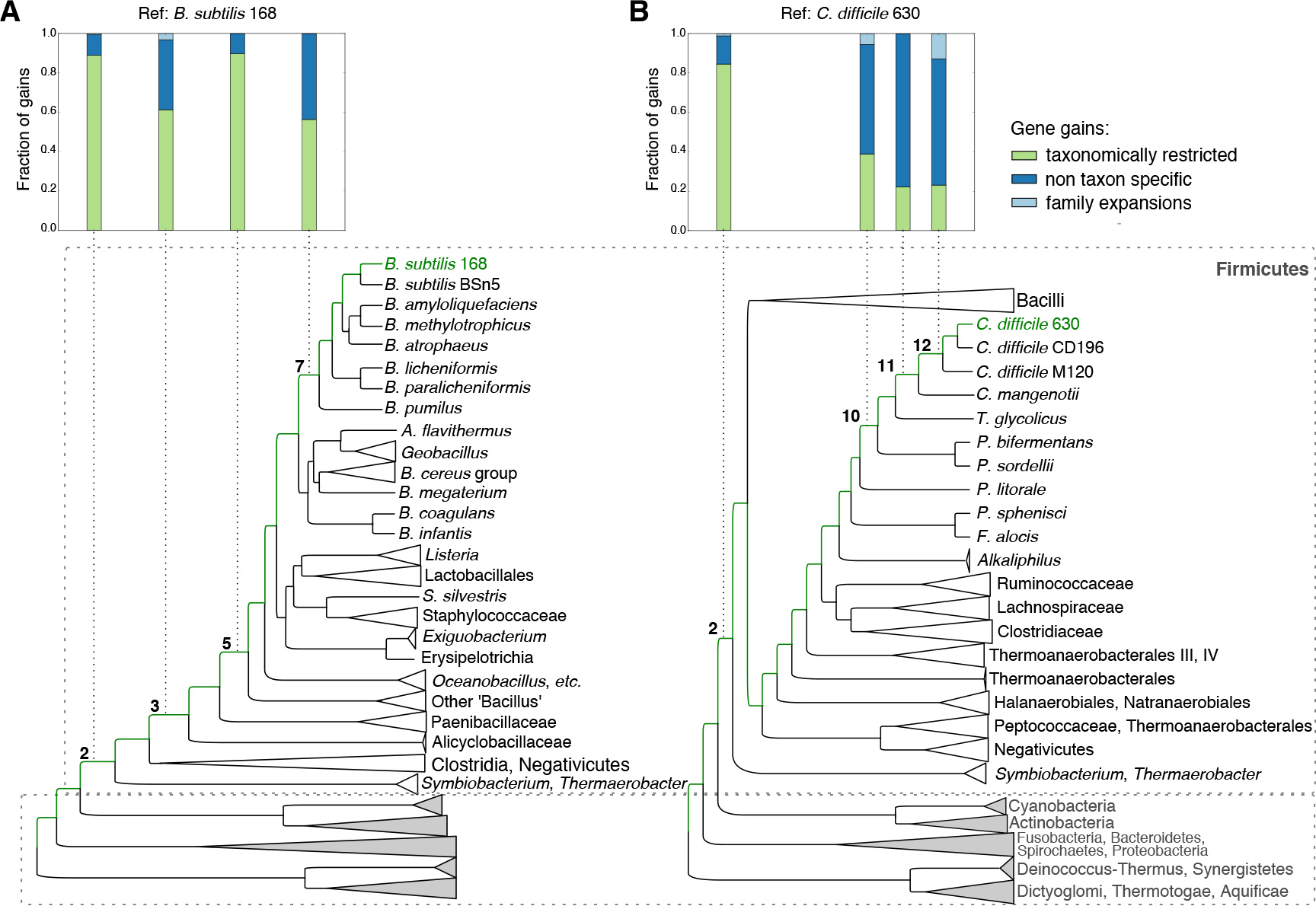
Predicted categories of sporulation gene gains in four branches for (A) *B. subtilis* and (B) *C. difficile.* Gene gains were considered to be taxon-specific (green) when gained at the branch, present in its descendants but not present or gained in other clades. Alternatively, gene gains were considered non-taxon specific (dark blue) when gained along a certain branch but also present in other non-descendent clades. The later included genes gained by HGT in both directions (from and to the branch/clade). Family expansions (light blue) were originated by duplication events.

### *B. subtilis versus C. difficile*: sporulation driven by rapid divergence and horizontal gene transfer

Besides the base of the Firmicutes we also estimated the origin of gene gains at lower taxonomic ranks. In particular we chose branches in the phylogenetic tree for which we could strictly define descendants and outgroups in agreement with the NCBI taxonomy (release from May 2018). These ranks, shown in Figure 5, included: branch 3 (at the base of the class Bacilli), branch 5 (Bacilli families but excluding the early diverging Alicyclobacillaceae and Paenibacillaceae), branch 7 (at the base of the *B. subtilis* group), branch 10 (within the family Peptostreptococcaceae - a subgroup containing the genera *Clostridioides*, *Paeniclostridium*, *Paraclostridium* and *Terrisporobacter*), branch 11 (at the base of the genus *Clostridioides*) and branch 12 (at the base of the species *C. difficile*).

At the base and within Bacilli, the estimated gains of TRGs were dominant over HGTs, representing 61,3% (branch 3), 89,8% (branch 5) and 56,3 % (branch 7) of the acquired genes (Figure 5A, in green). In contrast, genes acquired along the clostridial lineages were more often present in other clades (Figure 5B, in blue) and likely acquired by HGT, representing 55,6% in branch 10, 77,8% in branch 11 and 64,1% of the total gains in branch 12. Family expansions were significantly lower when compared to other mechanisms of gene acquisition with a maximum of four duplicated genes in branches 10 and 12. List of HGTs, TRGs and expansions by gene duplication are summarized in Tables S9 for Bacilli and S10 for Clostridia. Beyond the differences in rates of HGTs *vs.* TRGs and family expansion, we can find evidence of both mechanisms occurring in Bacilli and Clostridia (Figure 5).

### HGT and rapid divergence in Clostridia

In Clostridia we find CdeA, a protein involved in the exosporium assembly in *C. difficile* (Díaz-González et al. 2015; Calderón-Romero et al. 2018) coded by a gene gained in branch 10 (Entrez GI: 126699990, Table S10) which has orthologous outside the Peptostreptococcaceae family, notably in species of the intestinal tract (Figure 6). The orthologous sequences share in common two blocks of a four cysteine conserved domain with unknown function (InterPro Ac. Nr.: IPR011437) and together provide a clear example of HGT, likely to have occurred in the gut ecosystem.

**Fig 6.**
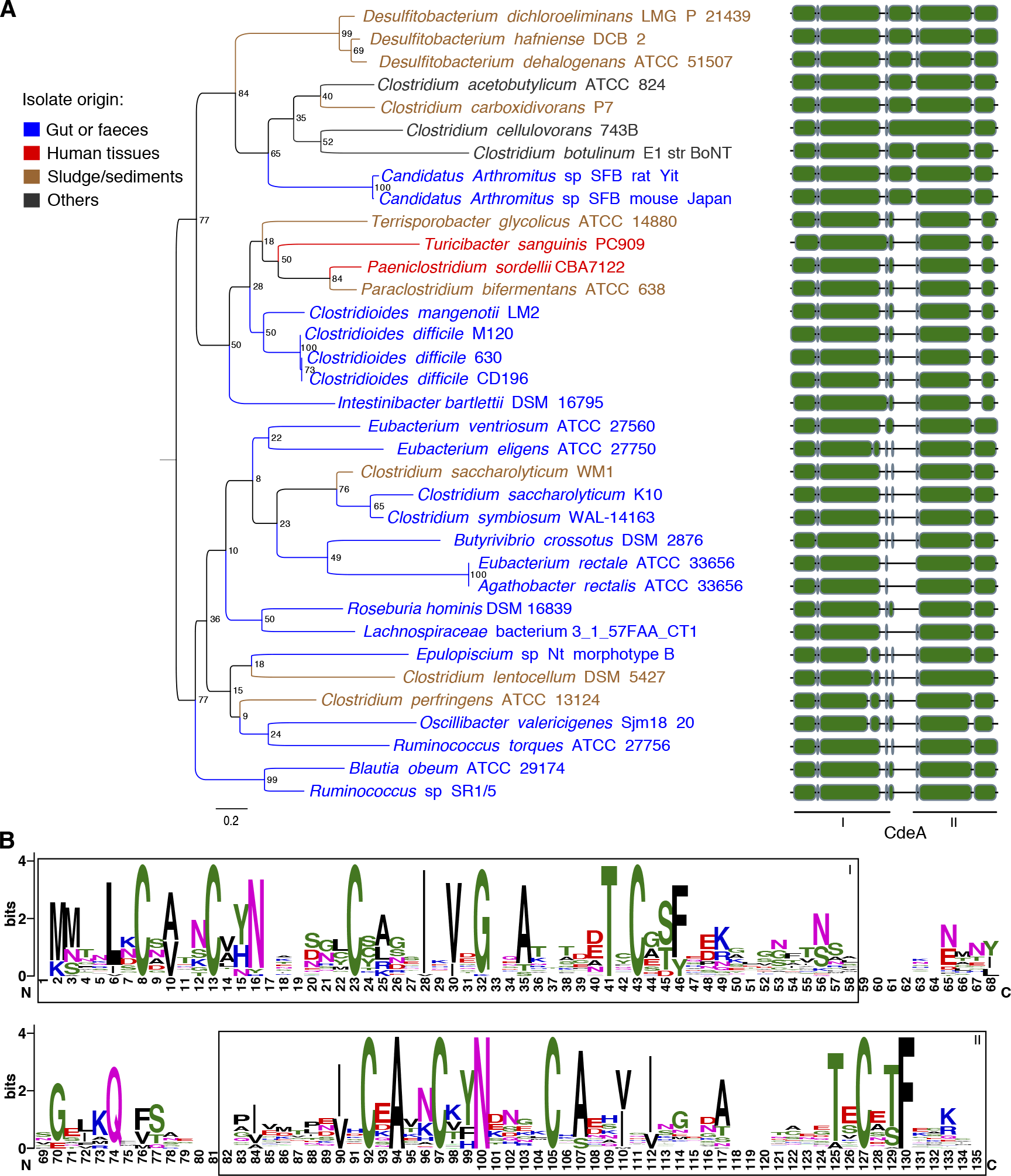
Phylogenetic tree and conserved blocks of CdeA proteins. Orthologous sequences were selected based on the results of the orthology mapping approach and supplemented with the orthologous cluster ENOG410XVF3 from EggNOG 4.5.1 (Huerta-Cepas, Szklarczyk, et al. 2016). Species are colored by environmental origin/niche. (A) Phylogeny with mid-point rooting and bootstrap probabilities were estimated with RAxML (Stamatakis 2014) using the best protein evolution model (LG+I) selected with ProtTest version 3 (Darriba et al. 2011), sequences were aligned using MAFFT (Katoh and Standley 2013). Protein blocks were drawn with ETE3 (Huerta-Cepas, Serra, et al. 2016). (B) Protein logos showing the two blocks of the conserved domain with four cysteines were estimated with WEBLOGO (Crooks et al. 2004).

In turn *spoiIIIAF* (Entrez GI: 126698793) has diverged to the point that no longer clusters with other genes that code for Stage III sporulation protein AF, forming two distinct orthologous clusters by our orthology mapping approach (Figure 7A) as well as in orthology databases EggNOG 4.5.1 and OMA: ENOG4105X2R and TLCVVIV for *spoiIIIAF*, ENOG410644B and IWEIGSE for *spoIIIAF* (Huerta-Cepas, Szklarczyk, et al. 2016; Altenhoff et al. 2018). The gene is part of the *spoIIIA* locus, conserved in all spore formers (Galperin et al. 2012; Morlot and Rodrigues 2018), which codes for eight proteins, SpoIIIIA to H, the components of a channel also known as the “feeding tube” and whose main role is to connect the mother cell to the forespore and transfer substrates required for σ^G^ activity in the forespore (Meisner et al. 2008; Camp and Losick 2009). Because *spoIIIAF* (Figure 7B) is poorly conserved, it is predicted here as an innovation ‘gained’ in branch 10, within the Peptostreptococcaceae family (Table S10). The *spoiIIIAF* cluster includes orthologous from all strains of *C. difficile*, *C. magenotii, Paraclostridium bifermentans, Paeniclostridium sordellii* and *Terrisporobacter glycolicus* (Figure 7A). An alignment of SpoIIIAF orthologs from both clusters on Figure 7B shows two-conserved TM domains (residues 1 - 57) and divergence in the remaining sequence, except for the hydrophobic residues highlighted in blue. These residues are part of the hydrophobic core of the protein which are associated with a conserved ring-building motif (RBM) fold (Zeytuni et al. 2018). Since a structure for SpoIIIAF is available (*B. subtilis*, PDB: 6dcs) (Zeytuni et al. 2018), we simulated the 3D structure of SpoiIIIAF from *C. difficile* using UCSF Chimera (Pettersen et al. 2004) and Modeller (Eswar et al. 2006) with 6dcs chain A as template. Simulations generated five reliable models (Figure S5 A), which are structurally similar to SpoIIIAF and adopt α1β1β2α2β4 topology (Figure 7C). Still, the high divergence rates between the two orthologs is reflected on the distribution of charged and hydrophobic residues on the protein surface (Figure 7C, Figure S5 B) and suggests that they have been under different evolutionary constraints.

**Fig 7.**
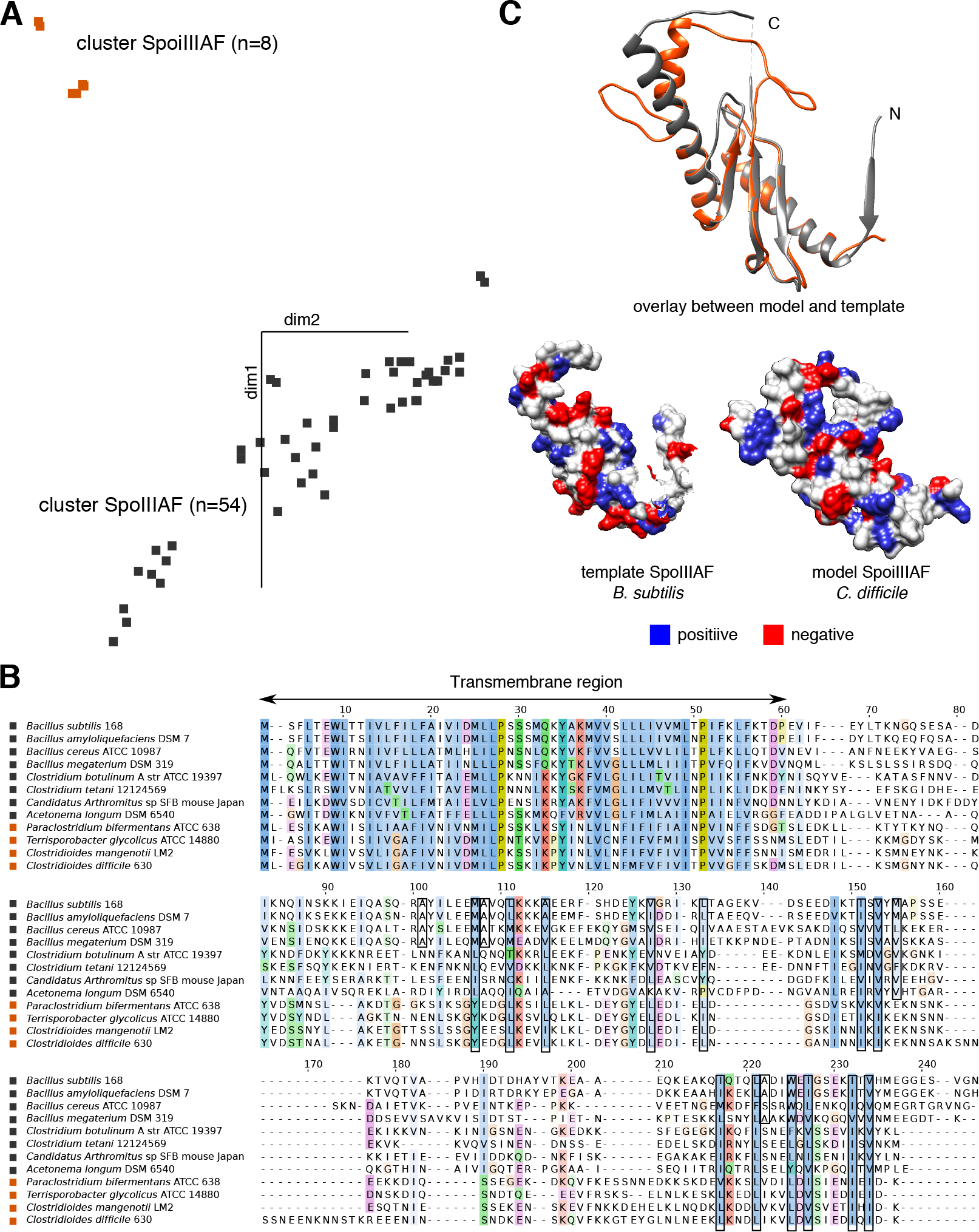
Comparative analysis between stage III sporulation proteins AF: SpoIIIAF from *B. subtilis* and SpoiIIIAF from *C. difficile*. (A) Orthologous clusters of SpoIIIAF and SpoiIIIAF (n=64) were merged and aligned using MAFFT (Katoh and Standley 2013) followed by a principal component analysis as implemented in Jalview (Waterhouse et al. 2009) to estimate sequence divergence between clusters. (B) Selected sequences clustering with SpoIIIAF (black square) and SpoiIIIAF (purple square) were aligned with T-coffee (Notredame et al. 2000). (C) Aligned structures of the monomeric forms of SpoIIIAF85-206 (grey) and SpoiIIIAF84-205 (orange) were determined using Modeller homology-based modeling (Eswar et al. 2006) with PDB:6dcs as template and MUSCLE (Edgar 2004) as sequence aligner. Structures and electrostatic charge distributions were visualized with Chimera (Pettersen et al. 2004).

## Discussion

In this work we have extended previous comparative genomics analyses based on *B. subtilis* sporulation genes (Paredes et al. 2005; de Hoon et al. 2010; Galperin et al. 2012; Abecasis et al. 2013; Galperin 2013) by including the sporulation genes identified in the human pathogen *Clostridioides difficile*. Recent advances in *C. difficile* genetics and the ability to construct sporulation mutants in this pathogenic anaerobe have enabled the identification of over 300 genes, directly regulated by the cell type-specific sigma factors of sporulation (Fimlaid et al. 2013; Saujet et al. 2013; Pishdadian et al. 2015) (Table S1). By comparing the sporulation-specific regulons between the two species we identified four main groups of genes based on their orthology/paralogy relationships. The largest group, representing over 50% of each regulon in both species, consist of genes only present in one species (Figure 1, Table S5). These genes highlight the existing diversity and complex origins of the sporulation machinery, genes coding for the regulatory complex SpoIVFA/SpoIVFB/BofA present in the forespore membrane of *B. subtilis* and involved in the activation of σ^K^ (Rudner 2002), are absent in the *C. difficile* genome (Pereira et al. 2013), as well as in other Clostridia species (Galperin et al. 2012). In some cases, genes having the same function in both species (*i.e.* functional equivalents) are annotated with the same name but have distinct evolutionary paths, with different ancestors, as is the case of the forespore proteins SpoIIQ. These proteins form a complex with SpoIIIAH through which the mother cell is thought to convey to the forespore metabolites essential for maintaining its metabolic potential and regulating engulfment in *B. subtilis* (Camp and Losick 2008; Meisner et al. 2008) and *C. difficile* (Fimlaid et al. 2015; Serrano et al. 2016). Despite having the same name and similar roles in both species, they are non-orthologous. Both SpoIIQ have a catalytic peptidase M23 motif (InterPro Ac. Nr.: IPR016047) and a transmembrane domain but distinct N-terminal sides, being a recognized case of non-orthologous gene displacement (Galperin et al. 2012). The second largest group of genes corresponds to those present in the two genomes but only described as part of the sporulation regulons in one species (Figure 1, Table S4). These genes are involved in other processes and may have been co-opted for roles in sporulation. For instance the genes *dapA* and *dapB*, which are present in many bacterial phyla and involved in the lysine biosynthesis pathway (Mukherjee et al. 2017), have been recruited to the sporulation process in *B. subtilis*, with *dapA* being part of the spoVF operon, expressed in the mother cell and involved in the synthesis of dipicolinic acid (DPA) (Chen et al. 1993; Daniel and Errington 1993; Steil et al. 2005), a compound that is characteristic of spores (Slieman and Nicholson 2001; Moeller et al. 2014). The same genes *dapA* and *dapB* in *C. difficile* are located in the *lysA-asd-dapA-dapB* operon (Mukherjee et al. 2017) and were shown to be dispensable for spore formation (Donnelly et al. 2016) whereas a study on *C. perfringens* suggests a more ancestral mechanism for DPA synthesis, involving the electron transfer flavoprotein Etfa (Orsburn et al. 2010). In the same group, the gene *uppS* (undecaprenyl pyrophosphate synthetase) is involved in polysaccharide biosynthesis and part of σ^G^ regulon in *C. difficile* (Saujet et al. 2013) but a role in *B. subtilis* sporulation has not been determined. Finally, there are two groups of orthologous genes in *B. subtilis* and *C. difficile*. One group corresponds to genes present in the same regulon in both species: 2 in σ^F^, 29 in σ^E^, 11 σ^G^ and 5 in σ^K^ (Table S2). For example the genes *spmA* and *spmB*, under the control of σ^E^ and involved in spore core dehydration (Popham et al. 1995; Steil et al. 2005; Pereira et al. 2013; Saujet et al. 2013). Other examples include the genes of the *spoIIIA* locus, also controlled by σ^E^ whose products are part of the SpoIIQ-SpoIIIAH channel (Illing and Errington 1991; Camp and Losick 2008; Fimlaid et al. 2015), and *spoIID* (σ^E^) required for septal peptidoglycan hydrolysis at the onset of the engulfment process and for the proper localization of SpoIIQ (Rodrigues et al. 2013; Serrano et al. 2016). In the σ^G^ regulon, the *pdaA* gene, coding for a N-acetylmuramic acid deacetylase, is involved in spore cortex peptidoglycan biosynthesis in both species (Fukushima et al. 2002; Saujet et al. 2013; Diaz et al. 2018). Still, despite being orthologs and regulated by the same sigma factors, these genes may show different modes of action in the two species through the link to non-homologous intermediates (Fimlaid et al. 2015; Serrano et al. 2016). At last, there are also orthologous pairs that belong to different regulons in the two organisms (Table S3). For example, the *seaA*, involved in the spore envelope assembly (Steil et al. 2005), is expressed early in the forespore, under the control of σ^F^ in *B. subtilis*, but is only activated following engulfment completion, under the direction of σ^G^ in *C. difficile* (Saujet et al. 2013). Also, *yraF,* coding for an uncharacterized protein similar to a spore coat protein, is transcribed in the forespore under the control of σ^G^ in *B. subtilis* (Wang et al. 2006), while in *C. difficile* is transcribed in the mother cell under the control of σ^E^ (Saujet et al. 2013).

We also find that some sporulation genes are conserved across several bacterial phyla, or even extended to other domains of life. For *e.g.* spoVR (orthologous group Nr.: COG2719) (Huerta-Cepas, Szklarczyk, et al. 2016) is present in Proteobacteria and Archaea. The gene *cotR*, with a conserved hydrolase domain (InterPro Ac. Nr.: IPR016035) and structure, has a broader taxonomic profile, which includes eukaryotes (orthologous group Nr.: COG3621) (Huerta-Cepas, Szklarczyk, et al. 2016). Still, genes of broader taxonomic range constitute less than one third of the sporulation machinery (Figure 3, Tables S9-S10). The great majority of sporulation genes were acquired in the first major gain event, coinciding with the origin of the Firmicutes (Figure 3) and including the genes of the so-called sporulation signatures based on *B. subtilis* genes (Abecasis et al. 2013) and spore-formers from the gut microbiota (Browne et al. 2016). Since the divergence of Firmicutes is estimated to have occurred between 2.5 to 3.0 billion years ago (Ga), upon the emergence of terrestrial ecosystems (2.6 Ga) and after the Great Oxidation Event (2.3 Ga) (Battistuzzi et al. 2004; Battistuzzi and Hedges 2009; Marin et al. 2017) it has been suggested that sporulation is part of an adaptive process to cope with O2 exposure (de Hoon et al. 2010; Abecasis et al. 2013). Gain events occurring after the origin of sporulation, and highlighted in Figure 3, correspond to lineage specializations of the program. The fact that we only find two larger gain events (≥90) along the evolutionary paths of *B. subtilis* and *C. difficile*, suggests that sporulation has evolved in punctual moments for these two species, only tied to certain lineages. Moreover, we find no evidence of these gains having greater influence in the size of a specific regulon since they have all co-evolved proportionally (Figure 4). This and the fact that the four sporulation-specific sigma factors were present in the last common ancestor of Firmicutes (Table S9, Table S10) suggest that both early and late sporulation stages were defined upon the emergence of the process.

Though the common macroevolutionary patterns in both lineages, our analysis to detect possible origins of gene gains, suggests different modes of gene acquisition. While in *B. subtilis* most of the genes gained in branches 3, 5 and 7 are taxon-specific (Figure 5A, Table S9), in the evolutionary path of *C. difficile*, genes were mostly acquired by horizontal gene transfer events, with a consistently smaller proportion of taxonomically restricted genes (Figure 5B). This finding is in line with the highly mobile genome of *C. difficile* 630, where 11% of mobile genetic elements (mainly transposons) are the main cause for acquisition of foreign genes from other species in the gut microbiota (Sebaihia et al. 2006; He et al. 2010). Genes acquired by HGT are involved in antibiotic resistance, virulence and sporulation, conferring *C. difficile* adaptive advantages in the gut ecosystem. A case-study emerging from our analysis is the *cdeA* gene, coding for an exosporium protein (Díaz-González et al. 2015) only found in Clostridia, especially in species inhabiting gut ecosystems (clusters ENOG410XVF3 and HESKKIA from EggNOG 4.5.1 and OMA databases). A phylogenetic tree based on the alignment of the CdeA sequences (Figure 6) does not reflect the species phylogeny suggesting multiple HGT events within the gut environment and through the cycling of clostridia species among different niches, *i.e.* mostly between sludge/sediments and the gut.

In turn *B. subtilis* shows a stronger lineage-specific specialization of the program, with genes that are only found in closely related species (Figure 5A, Table S9). The appearance of a higher number of such genes in members of the *Bacillus subtilis* group, may be due to the higher homologous recombination rates observed for Bacilli species when compared to *C. difficile* (Vos and Didelot 2009; He et al. 2010). A key feature of *B. subtilis* and close relatives is their natural competence, *i.e.* the ability to enter a physiological state in which cells are able to take up exogenous DNA fragments (Dubnau 1991; Haijema et al. 2001; Smits et al. 2005; Chen et al. 2007). Although the consequences of natural competence on the genomic composition and species diversity remain to be fully elucidated, this mechanism of DNA uptake as been proposed as a main driver of genome diversity within the *Bacillus subtilis* group (Brito et al. 2018) and to facilitate homologous recombination (Vos 2009; Mell and Redfield 2014).

The other mechanism giving rise to TRGs are high mutation rates that saturate the signal of homology. Rapid divergence between orthologs may occur driven by exposure to particular lineage-specific environment/niches and to different protein interactions/binding partners. This latter is illustrated here for the counterparts of SpoIIIAF from members of the Peptostreptococcaceae family (SpoiIIIAF) (Figure 7A). This protein is a component of the feeding tube, conserved in spore-formers, connecting the mother cell to the forespore, which together with other proteins of the spoIIIA operon: SpoIIIAB, SpoIIIAD, SpoIIIAE, SpoIIIAG form a multimeric membrane complex with a ring shape located in the mother cell membrane and extending to the intermembrane space between mother cell and forespore (Doan et al. 2009). Although the role of SpoIIIAF in the complex has not yet been fully elucidated, the protein contains a Ring-Building motif that may support ring formation in a multimeric form, similar to SpoIIIAG and SpoIIIAH (Levdikov et al. 2012; Zeytuni et al. 2017; Zeytuni et al. 2018). Like in *B. subtilis* and other spore-formers, SpoiIIIAF in *C. difficile* preserves the N-terminal transmembrane region and the RBM hydrophobic motif (Figure 7B). Still, the sequence as diverged considerably, only sharing 16% of sequence identity with its counterpart in *B. subtilis* (Serrano et al. 2016). This divergence is reflected in changes in the overall electrostatic potential and hydrophobicity on the proteins surface (Figure 7C, Figure S5). Since SpoIIIAF is predicted to be located in the inner mother cell membrane towards the cytoplasm (Doan et al. 2009; Zeytuni et al. 2017; Morlot and Rodrigues 2018), the high sequence divergence between SpoiIIIAF and SpoIIIAF suggests links with distinct proteins or binding partners in the mother cell cytoplasm that need to be further explored for Bacilli as well as for the members of the Peptostreptococcaceae family.

Besides gene-gains there were also extensive gene losses occurring independently throughout the bacterial phylogeny (Figure 3). These losses have resulted in the emergence of asporogenous lineages (*i.e.* non-spore-forming Firmicutes, (Onyenwoke et al. 2004)) among spore-formers at different taxonomic levels. Within Bacilli, we find the group including *Listeria*, Lactobacillales, Staphylococcaceae, *Solibacillus silvestris* StLB046 and *Exiguobacterium*; in Clostridia, the groups including the species *Proteocatella sphenisci* and *Filifactor alocis*, members of the family Lachnospiraceae and the genus Caldicellulosiruptor (within Thermoanaerobacterales group III) (Figure S1). Although the ability to form endospores is often seen has an evolutionary advantage, the process is energetically costly, involving the expression of hundreds of genes and taking up several hours to complete (Fujita and Losick 2005; Paredes et al. 2005; Fimlaid et al. 2013). For example, in nutrient-rich environments after 6,000 generations of *B. subtilis*, neutral processes but also selection were found to facilitate loss of sporulation (Maughan et al. 2007), occurring by indels or single-nucleotide substitutions (Maughan et al. 2009). In nature though, sporulation loss is mostly driven by consecutive gene losses as suggested earlier (Galperin et al. 2012; Galperin 2013) and demonstrated in Figure 3. Under relaxed environments for sporulation, emerging asporogenous bacteria will have less genes, be more energy efficient and grow faster (Maughan and Nicholson 2011; Nicholson 2012; Maitra and Dill 2015). Asporogenous Firmicutes have developed alternative ways to persist in diverse (Sauders et al. 2012; Linke et al. 2014) and sometimes extreme environments, as is the case of the extremophiles *Exiguobacterium* (Vishnivetskaya and Kathariou 2005; Vishnivetskaya et al. 2009; Moreno-Letelier et al. 2012) and *Caldicellulosiruptor* (Van De Werken et al. 2008; Hamilton-Brehm et al. 2010). The two *Exiguobacterium* species used here, *E. antarcticum* and *E. sibiricum,* have only 21% of the sporulation signature (Abecasis et al. 2013) while *Caldicellulosiruptor* species have between 62.5 and 73%, so there is still some speculation as to whether the latter form spores or not (Abecasis et al. 2013). The same question has persisted for sometime for *Ruminococcus* species (Galperin et al. 2012; Abecasis et al. 2013). *Ruminococcus* sp. have lost some of the conserved sporulation genes (Figure 3B) and have between 48 and 60% of the sporulation signature (Abecasis et al. 2013) (Figure S1, Figure S3), in what appears to be an intermediary stage into becoming asporogenous. Still, they were recently shown to sporulate (Browne et al. 2016; Mukhopadhya et al. 2018), and are likely to spread as spore-like structures between hosts (Schloss et al. 2014; Mukhopadhya et al. 2018), which plays as a counter evolutionary force to the emergence of asporogenous phenotypes. In other cases, notably *Epulopiscium* and *Candidatus Arthromitus*, sporulation has evolved as a reproductive mechanism, involving some gene loss (Figure 3B) and eventually the gain of new genes (Flint et al. 2005; Galperin 2013), under distinct evolutionary forces.

## Conclusions

In this study we were able to combine the comprehensive knowledge of sporulation genes in *B. subtilis* with the emerging model *C. difficile*. Our analysis validates earlier studies in that sporulation emerged once, at the base of Firmicutes. From this first major gene gain event, we have identified intermediate evolutionary steps combined with a second major lineage-specific gain event that underlies the current specialization and diversity observed among *B. subtilis* and *C. difficile* sporulation programs. Furthermore, predominant mechanisms of gene acquisition and innovation differ in the lineages of the two species: *B. subtilis* entails greater innovation with many TGRs while *C. difficile* has acquired new genes mostly through HGTs. We also show that extensive gene losses underlie the emergence of asporogenous lineages and the co-option of sporulation as a reproductive mechanism. Overall our comparative genomics approach provides a framework with which to elucidate the evolution and diversity observed in the sporulation program across Firmicutes and unveils new evolutionary case studies, such as those described for the genes *cdeA* and *spoIIIAF*.

## Methods

### Datasets construction

All known sporulation genes from *Bacillus subtilis* strain 168 and *Clostridioides difficile* strain 630, the two species that are better characterized in terms of their sporulation machinery, were collected from the literature (Fimlaid et al. 2013; Saujet et al. 2013; Pishdadian et al. 2015; Meeske et al. 2016) and SubtiWiki 2.0 (Mäder et al. 2012). These included the genes coding for the main regulators of sporulation: spo0A - controlling initiation; sigE and sigK - coding for the mother cell early and late specific sigma-factors; sigF and sigG - coding for the early and late forespore-specific sigma factors; and the genes under their control (Table S1). Bacterial genomes (n=258) were collected from the NCBI Reference Sequence Database (release from April 2017) including species from the classes Actinobacteria (15), Aquificae (2), Bacteroidetes (3), Cyanobacteria (3), Dictyogiomi (2), Firmicutes (186), Fusobacteria (4), Proteobacteria (30), Spirochaetes (6), Synergistetes (1), Thermotogae (5), Deinococcus-Thermus (1) (Figure S1, Table S6).

### Orthology/paralogy mapping and selection of homologous clusters

The predicted proteins from each genome were categorized into homologous clusters using the bidirectional best hits (BDBH) algorithm implemented in GET_HOMOLOGUES (Contreras-Moreira and Vinuesa 2013) (Figure S2). The software is built on top of BLAST+ (Camacho et al. 2009). First, inparalogues, defined as intraspecific BDBHs, are identified in each genome. Second, new genomes are incrementally compared to the reference genome and their BDBHs annotated. The main script ran with BLASTP default options and a minimum coverage fixed at 65% for the pairwise alignments (-C), option -s to save memory, and reporting at least clusters with two sequences (-t set at 2). The choice of the BDBH algorithm and corresponding cutoffs resulted from prior validation with sets and subsets of the bacterial genomes used in this study (Figure S3). The method was applied twice, the first time with *Bacillus subtilis* strain 168 (BSU) genome fixed as reference (-r) and the second time with *Clostridioides difficile* strain 630 (CD) (Figure S2). Homology mapping resulted in over 3500 clusters of homologous genes for references BSU and CD. Clusters corresponding to sporulation genes were selected making a total of 726 and 307 clusters for BSU and CD, respectively. The results were represented in two large-scale profile matrices of presence/absence of genes (0 for absence, 1 presence or >1 for multiple presence). The two reference genomes were also compared between each other using the orthoMCL algorithm implemented in GET_HOMOLOGUES. The main script ran with BLASTP default options, -C set at 65 % and reporting all clusters (-t 0).

### Bacterial phylogenies

Seventy gene clusters of single-copy orthologous conserved across the genomic dataset (adk, alaS, aspS, coaD, cysS, efp, engA, fabG, fmt, frr, gcp, gltX, guaA, infA, infC, ksgA, miaA, mraA, mraW, mraY, murC, murD, murE, murG, obgE, pheS, pheT, prfA, pth, pyrH, rluD, rpe, rplA-F, rplI-L, rplN-P, rplR-X, rpmA, rpoA, rpsB-E, rpsH, rpsJ, rpsK, rpsM, rpsQ, rpsS, ruvB, uppS, uvrB, yabC, yabD, ylbH, yyaF) were assembled with good occupancy; allowing for a maximum of 5 sequences missing per cluster. Clusters were aligned independently using MAFFT (Katoh and Standley 2013) or MUSCLE (Edgar 2004) under the default options. The best alignment per cluster was selected and trimmed using trimAl (Capella-Gutiérrez et al. 2009) with the -automated1 option. The 70 alignments were concatenated into a single super-alignment comprising 258 species/strains and 14401 amino-acid positions. The alignment was used to construct phylogenetic trees by maximum likelihood and Bayesian methods (Figure S4). Calculations and tree construction were performed with the RaxML 8 (Stamatakis 2014) (-N 100) and Mr. Bayes 3.2.5 (Ronquist et al. 2012) using the best protein evolution model (LG+G) selected with ProtTest version 3 (Darriba et al. 2011). Tree topologies generated with both methods were compared using ete-compare (Huerta-Cepas, Serra, et al. 2016) based on the normalized Robinson-Foulds distance (nRF). The RF scores are the total number of leaf bipartitions occurring in one but not in both trees. Since non-normalized measures of phylogenetic distance typically increase with the number of leaves in the tree, we used the normalized version, nRF, which divides RF by the maximum possible RF value for *n* leaves (maxRF). For topologically identical trees nRF will be 0. Our nRF of 0.03 (RF/maxRF = 16.0/512.0) indicates similar tree topologies.

### Phylogenetic profiles and evolutionary analysis of sporulation genes

Sporulation gene clusters were represented in two large-scale profile matrices of presence/absence, sized 726×258 and 307×258, for references BSU and CD. The profiles were combined with the bacterial phylogeny to estimate gain (0 → 1), loss (1 → 0) and duplication events (1 → 2 or more) in every tree branch with the software Count v 10.04 (Csurös 2010). First, rates were optimized using the gain–loss–duplication model implemented in Count, allowing different gain–loss and duplication–loss rates in all lineages and rate variation across gene families (1 to 4 gamma categories). One hundred rounds of optimization were computed. Next, a gene family history analysis was computed by posterior probabilities based on a phylogenetic birth-and-death model. Tables with the posterior probabilities based on the two profile matrices (Tables S7-S8) were extracted for further analysis with custom Python scripts to estimate total number of gene gains, losses and duplications (per branch), and gene presences (per node).

### Estimating origins of gene gains

To estimate the source of gene gains only events with a posterior probability of ≥ 0.5 by Count (Csurös 2010) were considered. Gene gains were classified as ‘taxon-specific’ (TRGs) when present at a given branch, present in its descendants but not present in other clades. Alternatively, they were considered ‘non-specific’ when gained at a certain branch, present in its descendants but also present in other clades of the phylogeny. The later included genes gained by HGT in both directions (from and to the branch/clade), since it is often challenging to identify the donor (Koonin et al. 2001), specially when dealing with large genomic sets. At last, a small fraction of sporulation genes was also gained by gene duplication and classified as ‘family expansions’.

Since our ancestral reconstruction was limited to 258 bacterial genomes we needed to validate genes classified as ‘taxon-specific’ in a broader taxonomic context. To do so, we performed searches against the NCBI NR database (released in January 2018, Cut-offs: E-value 10E-10, 75% query coverage with 50% identity) using BLASTP and checked for their best hits outside the clade and corresponding taxonomic distribution. While using this approach we were confronted with several false positives in the NCBI nr database that were removed from the BLASTP reports either because of misidentified species, for *e.g.* a genome assigned to the species *Varibaculum timonense*, from Actinobacteria (Guilhot et al. 2017), that is in fact a Clostridia, or because of recurring genomic contaminations of spore-formers in genome assemblies of other species as was shown for *Mycobacterium* (Ghosh et al. 2009; Traag et al. 2010). Genes gained in the braches 2, 3, 5, 7, 10, 11 and 12 and classified as ‘taxon-specific’, ‘non-specific’ or expansions are listed in Tables S9 and S10 with corresponding cluster names and accession numbers.

## Acknowledgements

We thank José Pereira Leal for advice with the bioinformatics methods and analyses and the Computational Genomics Laboratory for helpful discussions.

## Funding

This work was supported by FCT, “Fundação para a Ciência e a Tecnologia”, Portugal through grants SFRH/BPD/103171/2014 to PRS, PEst-OE/EQB/LA0004/2011 to AOH and by program IF (IF/00268/2013/CP1173/CT0006) to MS. This work was also financially supported by project LISBOA-01-0145-FEDER-007660 (“Microbiologia Molecular, Estrutural e Celular”) funded by FEDER funds through COMPETE2020 – “Programa Operacional Competitividade e Internacionalização”.

## Authors’ contributions

PRS designed the study, collected datasets, implemented the bioinformatics methods, performed analyses, interpreted the data and wrote the manuscript. MS provided the manually curated sets of sporulation genes and contributed with scientific input for the manuscript. AOH designed the study, interpreted the data and co-wrote the manuscript.

## Competing interests

The authors declare that they have no competing interests.

