## Supplementary File 1: Figure S1 for "From root to tips: sporulation evolution and specialization in *Bacillus subtilis* and the intestinal pathogen *Clostridioides difficile*"

Javascript must be enabled to view this page.

magnitude
score

 258
 .445736434108527
 258
 .445736434108527
 15
 0
 13
 0
 2
 0
 2
 0
 2
 0
 6
 0
 3
 0
 3
 0
 3
 0
 2
 0
 1
 0
 2
 0
 2
 0
 2
 0
 3
 0
 3
 0
 3
 0
 2
 0
 1
 0
 1
 0
 1
 0
 1
 0
 1
 0
 1
 0
 2
 0
 2
 0
 1
 0
 1
 0
 1
 0
 1
 0
 1
 0
 1
 0
 3
 0
 2
 0
 2
 0
 2
 0
 1
 0
 1
 0
 1
 0
 1
 0
 1
 0
 1
 0
 3
 0
 3
 0
 1
 0
 1
 0
 1
 0
 1
 0
 1
 0
 1
 0
 1
 0
 1
 0
 1
 0
 1
 0
 1
 0
 1
 0
 1
 0
 1
 0
 2
 0
 2
 0
 2
 0
 2
 0
 2
 0
 186
 .618279569892473
 77
 .532467532467532
 53
 .773584905660377
 2
 1
 1
 1
 1
 1
 30
 1
 1
 1
 1
 1
 22
 1
 4
 1
 1
 1
 1
 1
 2
 0
 2
 0
 3
 0
 3
 0
 8
 1
 3
 1
 4
 1
 1
 1
 1
 1
 1
 1
 1
 0
 1
 0
 6
 0
 1
 0
 5
 0
 24
 0
 1
 0
 1
 0
 2
 0
 2
 0
 5
 0
 3
 0
 1
 0
 1
 0
 5
 0
 3
 0
 2
 0
 5
 0
 3
 0
 1
 0
 1
 0
 6
 0
 2
 0
 4
 0
 97
 .721649484536082
 69
 .753623188405797
 17
 1
 2
 1
 2
 1
 13
 1
 1
 0
 1
 0
 1
 1
 1
 1
 1
 1
 1
 1
 9
 .111111111111111
 1
 0
 1
 0
 1
 1
 2
 0
 1
 0
 1
 0
 1
 0
 16
 1
 1
 1
 2
 1
 3
 1
 3
 1
 5
 1
 1
 1
 1
 1
 11
 .727272727272727
 1
 0
 1
 1
 1
 1
 4
 1
 1
 0
 1
 1
 9
 .555555555555556
 1
 1
 1
 0
 4
 1
 3
 0
 1
 1
 1
 1
 2
 1
 1
 1
 1
 1
 5
 .6
 3
 .333333333333333
 2
 0
 1
 1
 2
 1
 1
 1
 1
 1
 1
 1
 1
 1
 1
 1
 22
 .636363636363636
 10
 1
 1
 1
 1
 1
 1
 1
 1
 1
 1
 1
 1
 1
 4
 1
 11
 .272727272727273
 8
 0
 2
 1
 1
 1
 1
 1
 1
 1
 1
 0
 1
 0
 1
 0
 1
 0
 11
 .363636363636364
 11
 .363636363636364
 2
 0
 2
 0
 9
 .444444444444444
 1
 1
 2
 0
 1
 1
 2
 0
 1
 1
 1
 1
 1
 0
 4
 0
 4
 0
 4
 0
 2
 0
 2
 0
 2
 0
 1
 0
 1
 0
 30
 0
 5
 0
 5
 0
 3
 0
 3
 0
 1
 0
 1
 0
 1
 0
 1
 0
 5
 0
 3
 0
 1
 0
 1
 0
 2
 0
 2
 0
 1
 0
 1
 0
 1
 0
 1
 0
 1
 0
 1
 0
 5
 0
 2
 0
 2
 0
 2
 0
 1
 0
 1
 0
 1
 0
 2
 0
 1
 0
 1
 0
 1
 0
 1
 0
 2
 0
 2
 0
 1
 0
 1
 0
 1
 0
 1
 0
 13
 0
 4
 0
 4
 0
 1
 0
 1
 0
 1
 0
 1
 0
 1
 0
 1
 0
 1
 0
 1
 0
 1
 0
 1
 0
 3
 0
 3
 0
 3
 0
 3
 0
 3
 0
 1
 0
 1
 0
 1
 0
 1
 0
 1
 0
 1
 0
 6
 0
 6
 0
 3
 0
 3
 0
 3
 0
 1
 0
 1
 0
 1
 0
 2
 0
 2
 0
 2
 0
 1
 0
 1
 0
 1
 0
 1
 0
 1
 0
 5
 0
 5
 0
 1
 0
 1
 0
 1
 0
 4
 0
 1
 0
 1
 0
 3
 0
 3
 0
