## Supplementary File 2: Figures S2 to S5 for "From root to tips: sporulation evolution and specialization in *Bacillus subtilis* and the intestinal pathogen *Clostridioides difficile*"

**Legends of Supplementary Figures in Supplementary Files 1 and 2:**

**Fig.S1. Taxonomic visualization of species sampled from the NCBI database (n=258).** Taxonomic ranks according to NCBI, (release from May 2018). Species and groups were classified as sporulating (in green) or non-sporulating (in red) based on the presence/absence of a sporulation genomic signature of 50 genes (cutoff $\geq$ 75%) (Abecasis et al. 2013). Results were displayed with Krona (Ondov et al. 2011).

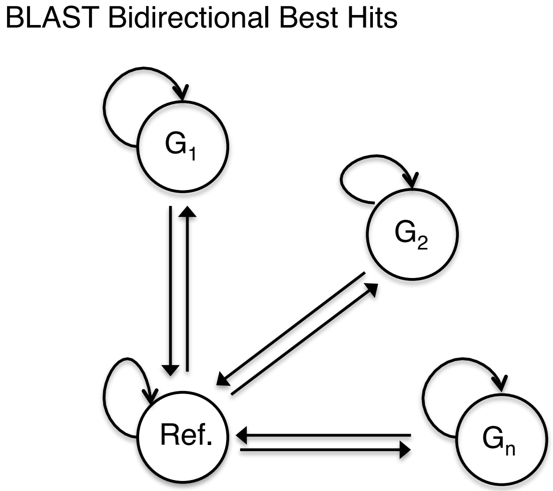


Ref. #1 *Bacillus subtilis* 168
Ref. #2 *Clostridioides difficile* 630

**Fig.S2. Schematic view of the homology mapping approach** using the algorithm BDBH as implemented in the software GET_HOMOLOGUES (Contreras-Moreira and Vinuesa 2013): first, inparalogues, defined as intra-specific bidirectional best hits (BDBHs), are identified in each genome; second, new genomes are incrementally compared to the reference genome and their BDBHs annotated; finally, clusters containing at least 1 sequence per genome are conserved. The algorithm was applied twice. First using the genome of *Bacillus subtilis* 168 as reference, second using the genome of *Clostridioides difficile* 630.

**A**

| Algorithm | Nr. of clusters | % sporulation clusters | TPR | FPR |
|---|---|---|---|---|
| BDBH | 3867 | 98 | 0,71 | 0 |
| OrthoMCL | 31733 | 100 | 0,81 | 0,42 |
| COGtriangles | 33162 | 100 | 0,79 | 0,88 |
| **Nr. of genomes** | | | | **42** |

**B**

| **% Signature** | **BBH coverage - 50%** | | | **BBH coverage - 65%** | | | **BBH coverage - 75%** | | |
|---|---|---|---|---|---|---|---|---|---|
| | FPR | TPR | ACC | FPR | TPR | ACC | FPR | TPR | ACC |
| **50** | 0,047 | 1,000 | 0,963 | 0,039 | 0,993 | 0,966 | 0,039 | 0,993 | 0,966 |
| **55** | 0,043 | 0,993 | 0,963 | 0,039 | 0,993 | 0,966 | 0,039 | 0,986 | 0,963 |
| **60** | 0,039 | 0,993 | 0,966 | 0,039 | 0,986 | 0,963 | 0,027 | 0,986 | 0,970 |
| **65** | 0,039 | 0,986 | 0,963 | 0,039 | 0,986 | 0,968 | 0,016 | 0,986 | 0,978 |
| **70** | 0,035 | 0,986 | 0,966 | 0,019 | 0,986 | 0,975 | 0,000 | 0,959 | 0,978 |
| **75** | 0,031 | 0,986 | 0,968 | 0,000 | 0,966 | **0,980** | 0,000 | 0,945 | 0,973 |
| **80** | 0,008 | 0,959 | 0,973 | 0,000 | 0,918 | 0,963 | 0,000 | 0,849 | 0,938 |
| **85** | 0,000 | 0,938 | 0,970 | 0,000 | 0,822 | 0,929 | 0,000 | 0,719 | 0,892 |
| **90** | 0,000 | 0,836 | 0,933 | 0,000 | 0,637 | 0,862 | 0,000 | 0,473 | 0,803 |
| **95** | 0,000 | 0,630 | 0,860 | 0,000 | 0,466 | 0,800 | 0,000 | 0,329 | 0,751 |
| **100** | 0,000 | 0,267 | 0,729 | 0,000 | 0,144 | 0,685 | 0,000 | 0,075 | 0,660 |
| **Nr. of genomes** | | | | | | | | | **405** |

**C**

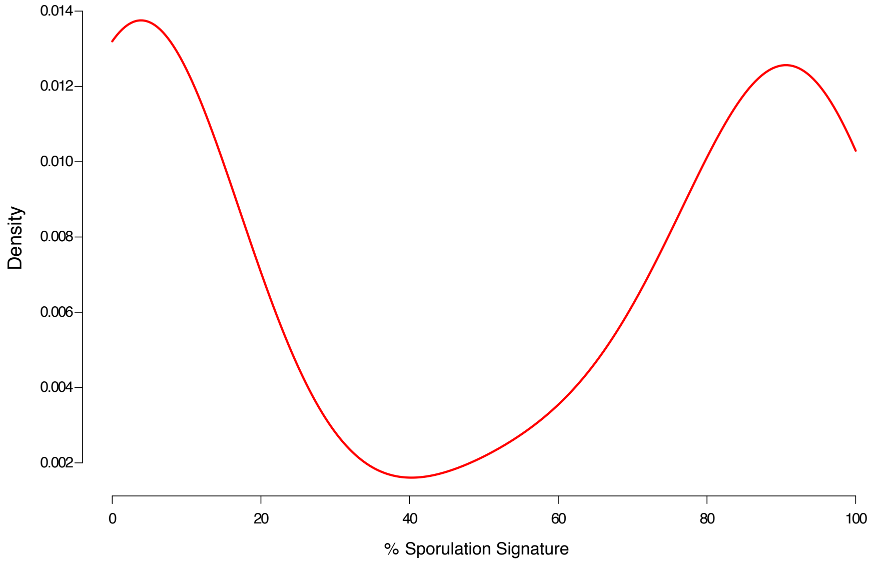

**Fig.S3. Validation of the homology mapping approach.** In order to choose the best algorithm and thresholds for orthology/paralogy mapping of sporulation gene families we selected sets of bacterial genomes, on which we had prior knowledge of their phenotype (*i.e.* spore forming or non-spore forming). Based on these sets it was possible to evaluate how good are the homology mapping algorithms (and thresholds) implemented in the GET_HOMOLOGUES software (Contreras-Moreira and Vinuesa 2013) at predicting true spore-formers and select the best approach for this study. Based on the presence/absence of a sporulation genomic signature of 50 genes (cutoff $\geq$ 75%) (Abecasis et al. 2013) we determined the false positive (FPR), true positive rates (TPR) and accuracy (ACC) at finding spore-formers between: (A) Different algorithms: BDBH, OrthoMCL and COGtriangles and (B) Different values of min coverage in BLAST pairwise alignments (option –C from GET_HOMOLOGUES) using the BDBH algorithm. We concluded that the best approach for orthology/paralogy mapping of sporulation genes would be to use the BDBH algorithm with option –C 65 and confirmed that a presence of at least 75% of the sporulation signature (Abecasis et al. 2013) is the best cutoff to classify a genome as belonging to a spore-former or not. (C) Distribution of the percentage of the sporulation signature present in the 258 bacterial genomes used in this work based on BDBH – coverage 65%. Spore-formers have high percentages of the signature (> 75%) while non-sporulating have less than 20%. Only few species (*e.g. Ruminococcus albus* DSM 20455, *Oscillibacter valericigenes* Sjm18-20, *Caldicellulosiruptor kristjanssonii* I77R1B, *Solibacillus silvestris* StLB046) have intermediate proportions of the signature, between 40 and 70%.

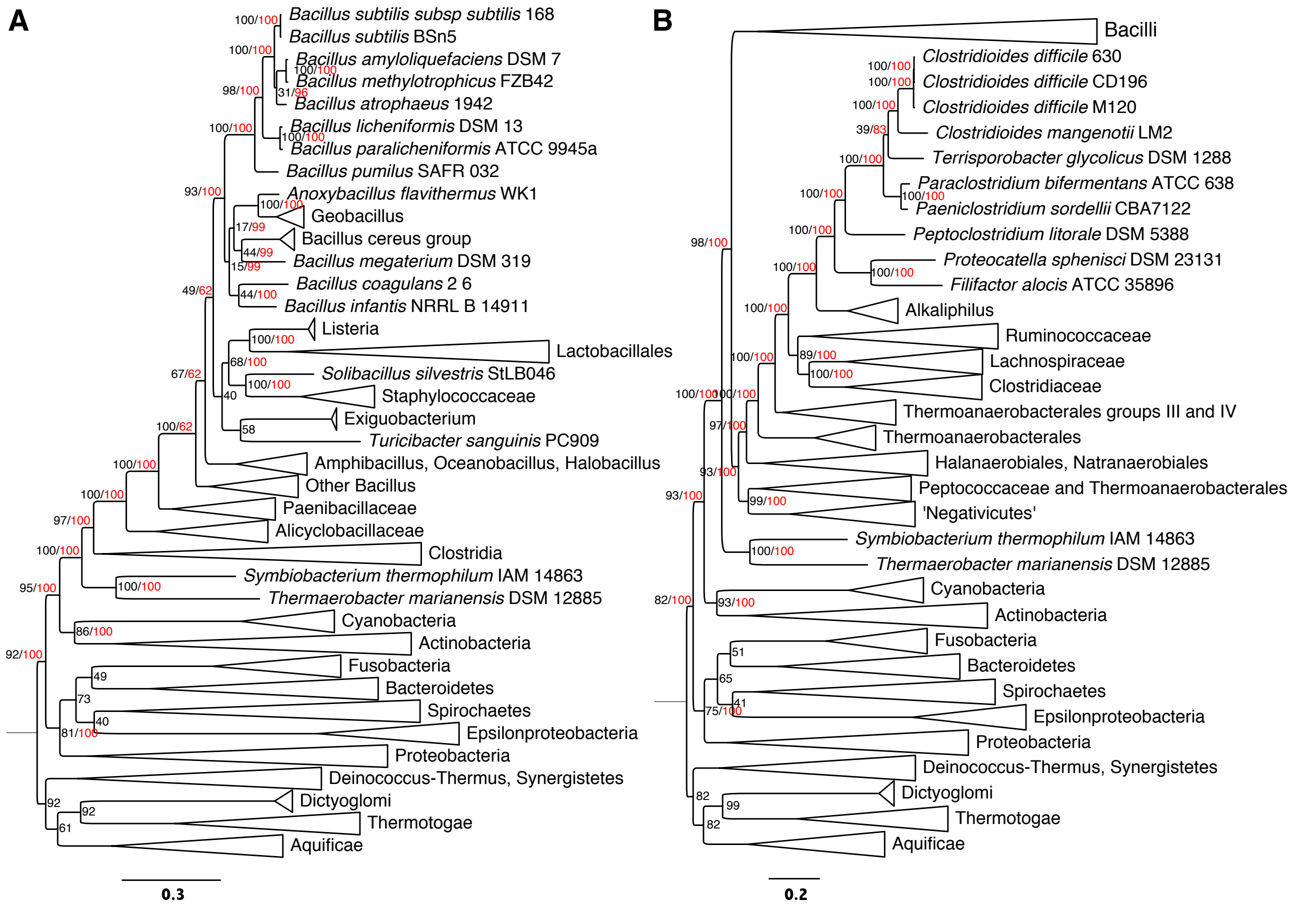


**Fig.S4. Bacterial phylogenies** highlighting main lineages of (A) Bacilli and (B) Clostridia with bootstraps measures from maximum likelihood (black) and Bayesian (red) methods implemented with RAxML (Stamatakis 2014) and Mr. Bayes (Ronquist et al. 2012).

**A**

| Model | zDOPE | Estimated RMSD | Estimated Overlap (3.5 ) |
|---|---|---|---|
| #1.1 | -0.50 | 6.772 | 0,447 |
| #1.2 | -0.50 | 6.631 | 0,452 |
| #1.3 | -0-54 | 6.728 | 0.453 |
| #1.4 | -0.66 | 6.945 | 0.425 |
| #1.5 | -0.62 | 7.723 | 0.349 |

**B**

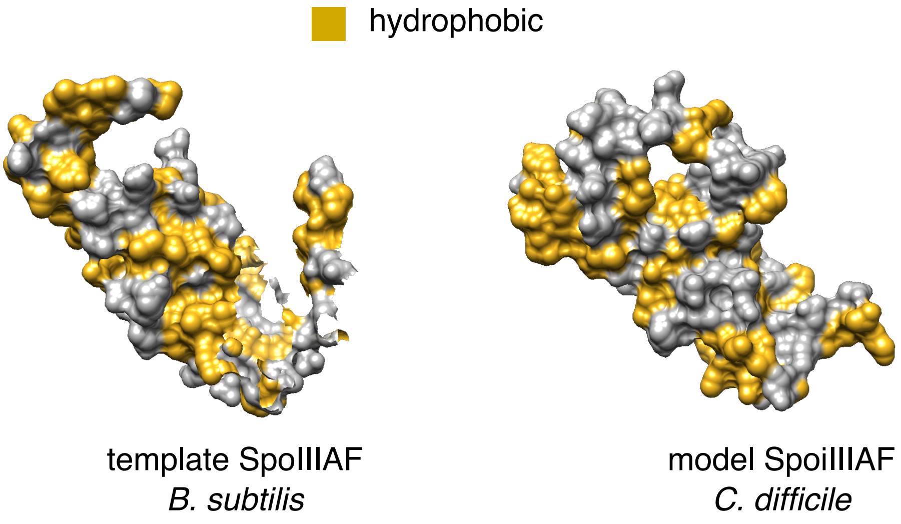


**Fig.S5. Comparative modeling approach with SpoIIIAF from *C. difficile* as model and SpoIIIAF from *B. subtilis* as template.** (A) Evaluation scores for the five models obtained with Modeller (Eswar et al. 2006) via Chimera interface (Pettersen et al. 2004). (B) Template and Model #1.3 were selected to map charged (red, blue in Figure 6 C) and hydrophobic residues on the protein surface (yellow).

**Legends of Supplementary Tables in Supplementary File 3:**

**Tab.S1. NCBI GenInfo Identifiers of sporulation genes** compiled from *B. subtilis* 168 (BSU) and *C. difficile* 630 (CD) with corresponding regulators.

**Tabs.S2-5. Results from the homology mapping between *B. subtilis* 168 and *C. difficile* 630**: sporulation genes with orthologous in both genomes and in the same regulon **(S2)**, sporulation genes with orthologous in both genomes but in different regulons **(S3)**, sporulation genes with orthologous in both genomes but only involved in sporulation in one species **(S4)**, sporulation genes without orthologous in the two species **(S5)**.

**Tab.S6. Bacterial genomes downloaded from NCBI** (release from April 2017) with corresponding GIs and Accession Numbers for closed genomes and draft (contig) genomes.

**Tabs.S7-8. Evolutionary analyzes of sporulation genes from *B. subtilis* 168 (S7) and *C. difficile* 630 (S8).** Posterior probabilities of presence, multi-members, gain, loss, expansion and contraction were computed with Count (Csurös 2010) for each sporulation gene family across the bacterial phylogeny. Total sums of posteriors per node (presences) and branch (gains, losses and duplications) were used to obtain expectations of total presences, gains and losses.

**Tabs.S9-10. Lists of sporulation genes from *B. subtilis* 168 present at the origin and gained at branches**: 2, 3, 5 and 7 **(S9)** and **Lists of sporulation genes from *C. difficile* 630 present at the origin and gained at branches**: 2, 10, 11 and 12 **(S10)** as predicted with phylogenetic birth-and-death model implemented with Count v 10.04 (Csurös 2010) using a threshold of 0.5 posterior probability. Gene gains were considered 'taxon-specific' when gained at the branch, present in its descendants but not present or gained in other clades. Alternatively gains were considered 'non-specific' when gained along a certain branch but also present in other non-descendent clades. Some gene gains initially predicted as taxon-specific were reclassified as 'non-specific' after validating with BLASTP against the NCBI NR database (release January 2018, Cut-offs: E-value 10E-10, > 75% query coverage, > 50% identity, frequency of hits outside the clade ≥ 25% - defined using specific taxonomic filters). Note: some gene gains may appear twice if posterior probability > 0.5, in which case one of the events will be more likely to occur.
